# Free-Standing Multilayer Films as Growth Factor Reservoirs for Future Wound Dressing Applications

**DOI:** 10.1101/2022.07.19.500607

**Authors:** Adrian Hautmann, Devaki Kedilaya, Sanja Stojanović, Milena Radenković, Christian K. Marx, Stevo Najman, Markus Pietzsch, João F. Mano, Thomas Groth

## Abstract

Chronic skin wounds place a high burden on patients and health care systems. The use of angiogenic and mitogenic growth factors (GF) can facilitate the healing but GF are quickly inactivated by the wound environment if added exogenously. Here, free-standing multilayer films (FSF) are fabricated from chitosan (CHI) and alginate (ALG) as opposing polyelectrolytes in an alternating manner using layer-by-layer technique (LbL). One hundred bilayers form an about 450 µm thick, detachable free-standing film (N-FSF) that is subsequently crosslinked by either ethyl (dimethylaminopropyl) carbodiimide (EDC) combined with N-hydroxysuccinimide (NHS) (E-FSF) or genipin (G-FSF). The characterization of swelling, oxygen permeability and crosslinking density shows reduced swelling and oxygen permeability for both crosslinked films compared to N-FSF. Loading of fibroblast growth factor 2 (FGF2) into the films results in a sustained release of GF from crosslinked in comparison to N-FSF. Biocompatibility studies *in vitro* with human dermal fibroblasts (HDF) cultured underneath the films demonstrate increased cell growth and cell migration for all films with and without FGF2. Especially G-FSF loaded with FGF2 greatly increases cell proliferation and migration. *In vivo* biocompatibility studies by subcutaneous implantation in mice show that E-FSF causes a strong inflammatory response while G-FSF is of high biocompatibility. N-FSF also represents a biocompatible film but shows early degradation. All FSF possess antibacterial properties against gram+ and gram-bacteria demonstrated by an agar diffusion disc assay. In summary, FSF made of ALG and CHI crosslinked with genipin can act as a reservoir for the sustained release of FGF2, possessing high biocompatibility in vitro and in vivo. Moreover, G-FSF promotes growth and migration of HDF and has antibacterial properties which makes it an interesting candidate for bioactive wound dressings.

## 1 Introduction

Chronic wounds represent a major problem in medical care, intensified by an ageing society. With a prevalence of about one percent in Germany (2012) it places a high burden on the quality of life of patients as well as the health care system [1]. Chronic wounds are non-healing wounds, which develop due to defective regulation of a tightly controlled healing process, often because of severe traumata and/or side effects from diseases, such as diabetes, age of patients and others [2]. Problems associated with chronic wounds, are extracellular matrix destruction, low amounts of oxygen (hypoxia), high amounts of reactive oxygen species and bacterial invasion. This results in a further progression of the wound, which can ultimately lead to sepsis and amputation of extremities associated with high mortality [2]. Very often these wounds are full thickness wounds; the epidermis, dermis and underlying structures are destroyed [3]. These kinds of wounds close by epithelial resurfacing and wound contraction. The normal wound healing process is characterized by four timely and locally orchestrated processes of hemostasis; inflammation, cell proliferation (granulation and reepithelization) and tissue remodeling [2]. The formation of granulation tissue for improved epithelial migration has been identified to be pivotal for the healing of chronic wounds [4]; The anti-inflammatory signals of macrophages shifted to an M2 state allows fibroblasts and keratinocytes to proliferate and migrate from the borders of the wound [4]. After the formation of granulation tissue, keratinocytes use this newly formed structure to migrate on top of the fibroblasts to reepithelize the wound [5].

To treat chronic wounds, a wide selection of advanced wound dressings has been developed such as semipermeable films, foam dressings, alginate (ALG) dressings, hydrogel, hydrocolloid, and hydrofiber dressings [6]. Their individual use case, advantages and disadvantages are discussed in a recent review published by Han et al.[6]. All wound dressings must be non-toxic, maintaining a moist wound environment while retaining the ability to absorb wound exudates, permitting gas exchange, and preventing bacterial invasion. Additionally, during the granulation phase, they should be non-adherent to the wound area [7]. To improve the resolution of the inflammation phase in chronic wounds, growth factor supplementation has gained particular attention since chronic wounds have been found to possess low amounts of growth factors, like platelet-derived growth factor (PDGF), epidermal growth factor (EGF), fibroblast growth factor (FGF), transforming growth factor (TGF-β) and vascular endothelial growth factor (VEGF) [8]. FGF2, also known as basic FGF (bFGF), has gained particular attention in recent years [9]. It plays an important role in the wound healing process, mainly by stimulating migration, proliferation, and differentiation of fibroblasts and keratinocytes cells, but also because of its angiogenic effects [9]. The application of FGF2 has shown positives outcomes in clinical studies of eardrum reconstruction, in periodontal regeneration, treatment of burns and diabetic ulcers [10–12]. However, FGF2 loses quickly its bioactivity in a wound due to degradation by proteases and other processes [13,14]. This requires the protection of growth factors (GF) by carrier systems for controlled release as inherent additional function of advanced wound dressings. FGF2-containing wound dressings were examined in several pre-clinical studies [15–17]. While the aforementioned FGF2 loaded wound dressings show successful application and efficacy of FGF2, these designs are hampered in either one of the following categories; they are monolithic in design and therefore limited in their choice of tuneable parameters, need demanding production requirements or possess lacking release kinetics of GF and other bioactive substances.

One technique interesting for wound healing applications is the layer-by-layer (LbL) method, based on the consecutive adsorption of oppositely charged polyelectrolytes on a substrate [18]. Biological building blocks, representing polyelectrolytes interesting for the area of wound dressings, are charged polysaccharides like ALG and chitosan (CHI), but also proteins like collagens or elastin [19]. In addition, particulate charged matter like microgels and nanoparticles that can be loaded with bioactive substances for medical applications represent building blocks for LbL, too [20,21]. LbL can be used for polyelectrolyte multilayer (PEM) coating of implants, scaffolds for tissue engineering applications or preparation of multilayer capsules used for cell immobilization or drug delivery applications [22]. In recent years the application of the technique was extended to the construction of detachable free-standing PEM films [23]. Such films can be prepared to a thickness in mm scale enabled by automatic dip or spray coating processes [24]. The substrate supported PEM films are freed either by dissolution of a sacrificial layer or by peeling off leading to the formation of free-standing films (FSF) [25]. Advantages of LbL technique as bottom-up process are that it is simple, cost effective and allows great customizability [24]. PEMs possess high loading capacities for biomolecules, drugs or metal ions and enable release time spans from weeks to months [26,27]. Based on these assets, bioactive functions, like anti-bacterial effects or immunomodulation of multilayer films were already realized [28,29]. Compared to other types of conventional wound dressings, the most relevant advantage of PEM-based FSF lies in their potential to be built in a modular manner. It allows the combination of different polyelectrolytes, bioactive substances, or particulate matter together or in separate sections of the film. Hence, a multitude of wound dressing related functions could be realized such as control of swelling and permeability, promotion of wound healing, as well as anti-oxidative and antibacterial functions.

Here, we were interested to use negatively charged ALG and positively charged CHI as building blocks for biogenic FSF prepared by LbL technique. ALG and CHI are both from biological origin, widely available, cost efficient and used as single components in several conventional wound dressings [30]. ALG in particular is valued for its ability to absorb water and body fluids [30]. CHI has antibacterial as well as anti-inflammatory effects and acts as a hemostatic agent [31]. Since GF like FGF2 can have a promoting effect on regeneration of wound tissue including neovascularization, we studied here the ability of FSF on uptake and release of FGF2 in dependence on crosslinking with either 1-ethyl-3-(−3-dimethylaminopropyl) carbodiimide (EDC) and *N*-hydroxysuccinimide (NHS) as well as genipin. Subsequently, the *in vitro* effects of these FSF on growth and migration of fibroblasts were studied as important prerequisites for regeneration of dermis. Since biocompatibility of such films is required for any clinical applications, we studied this in a mouse subcutaneous implantation model. Moreover, antibacterial activity of films was investigated using gram+ and gram-bacterial, which is another important aspect for the suggested use of these films. Results indicate that FSF crosslinked with genipin have a delayed release of FGF2, which promotes growth and migration of fibroblasts greatly, possess excellent biocompatibility in vivo and antibacterial activity, which makes this FSF highly interesting for wound healing applications. Results are reported herein.

## 2 Material and methods

### 2.1 Chemicals

Chitosan 85/500 with a deacetylation degree of 85% (CHI, Mw ≈ 500 kDa) was delivered from Heppe Medical Chitosan GmbH (Halle, Germany) and sodium alginic acid (low viscosity) from Alfa Aesar (ThermoFisher GmbH, Schwerte, Germany). Sodium chloride, acetic acid and *hydrochloric acid* were obtained from Carl Roth GmbH + Co. KG (Karlsruhe, Germany),

### 2.2 Cell culture

DMEM (without pyruvate, with 4.5 g/L glucose) (Carl Roth) supplemented with 10% (v/v) fetal bovine serum (FBS, Biochrom AG, Berlin, Germany) and 1% (v/v) antibiotic–antimycotic solution (AAS, Lonza, Basel, Switzerland) was used for culturing human dermal fibroblasts (HDF, PromoCell GmbH, Heidelberg, Germany) at 37 °C in a humidified 5% CO_2_/95% air atmosphere in a NUAIRE® DH Autoflow incubator (NuAire, Plymouth, Minnesota, USA). Cells were passaged every second day to maintain a maximum of 80% confluence in a T75 cell culture flasks (Greiner Bio-One GmbH and Co.KG, Frickenhausen, Germany). Trypsin/EDTA solution (0.25 % /0.02 % (w/v)) (BioChrom AG) was used to detach the adherent fibroblasts.

### 2.3 Multilayer formation

0.15 M NaCl solution (buffer and washing solution for dip coating), 2 mg/mL CHI solution and 5 mg/mL ALG solution (polymer reservoirs for dip coating) were prepared as previously described. [32]. The multilayered films were fabricated by the alternating deposition of CHI and ALG with intermediate washing steps using NaCl solution with an automated dip coater (DR01, Riegler & Kirstein, Berlin, Germany) to 100 double layers. The first layer was CHI and last layer ALG. Glass substrates were used for layer deposition (Menzel-Gläser, 76×26 mm^2^, Thermo Scientific, Hungary). Coating and washing times for every layer were 5 and 2.5 min, respectively. After dip coating, samples were washed with milli-Q H_2_O, manually detached and either crosslinked or stored at 4 °C for further experiments.

### 2.4 Film crosslinking

After detachment, films were punched into circular 12 mm diameter disks and placed inside 24-well plates. Two different crosslinking methods were applied, which mechanisms are illustrated in Figure 1. In the first case, films were immersed inside 1 mg/mL in genipin (Wako Chemicals GmbH, Neuss, Germany) in MilliQ water at 37 °C for 24 h. Absolute EtOH (Ethanol absolute, AppliChem Panreac ITW Companies, Darmstadt, Germany) was used to stop the reaction, followed by three 15 min washing cycles with Milli-Q water. Alternatively, crosslinking using 1-ethyl-3-(3-dimethylaminopropyl) carbodiimide hydrochloride (EDC) (Carl Roth GmbH + Co. KG) and N-hydroxysuccinimide (NHS) (Sigma-Aldrich Chemie GmbH, Steinheim, Germany) was applied. EDC was used at a concentration of 50 mg/mL, dissolved in 150 mM NaCl (pH 5.0) followed by the addition of NHS at 11 mg/mL in 150 mM NaCl. The crosslinking solution was added to the films that were subsequently incubated at +4 °C for 18 h. 0.15 M NaCl solution at pH 8.0 was used to remove the crosslinkers, washing the films for 1 h, repeated three times.

**Figure 1.**
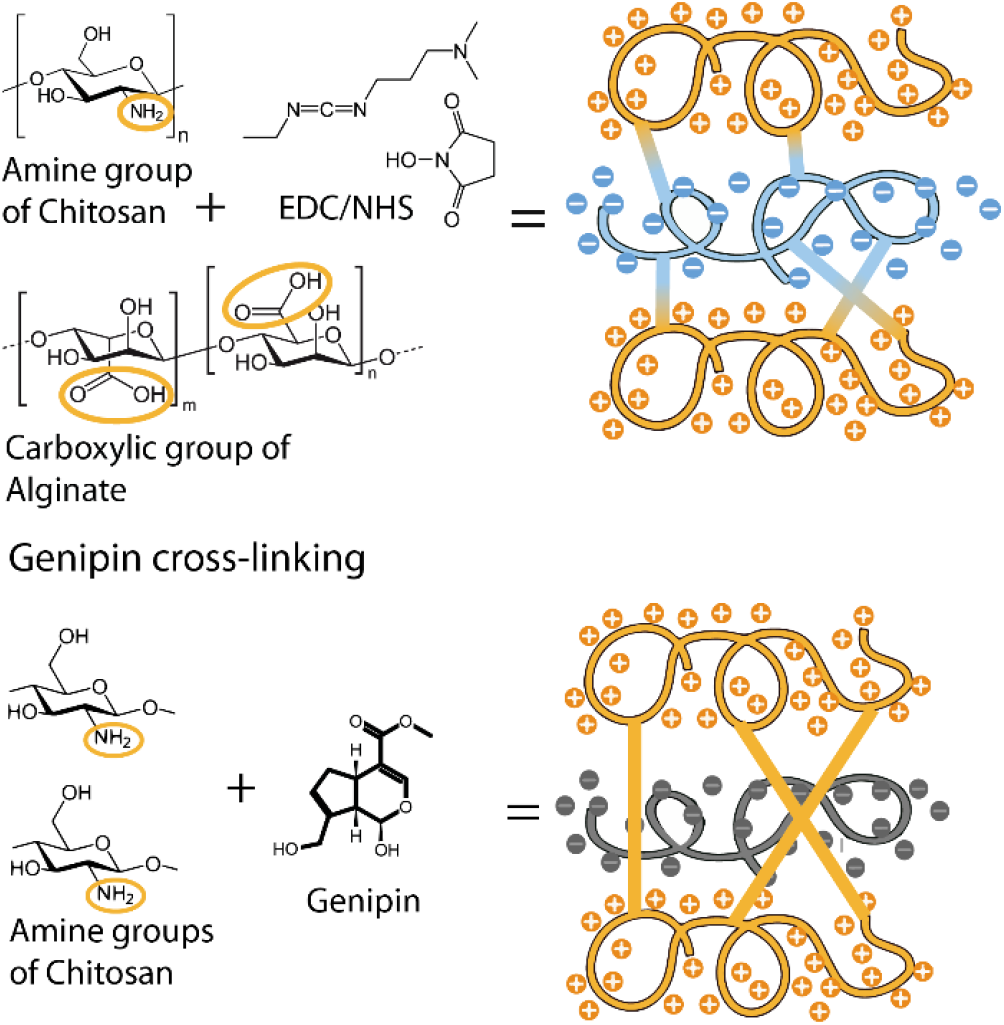
Crosslinking Chemistry – EDC/NHS: the carbodiimide conjugation works via the activation of carboxyl groups for direct reaction with primary amines via amide bond formation, which leads to the crosslinking of ALG with CHI. Genipin: The ester group of genipin reacts with amine groups of biomolecules forming an amide bond. In the case of the multilayer films, genipin crosslinks only CHI. The alginate layers are not crosslinked due to the absence of amine groups.

### 2.5 Determination of crosslinking degree

The crosslinked films were dried in a desiccator for 72 h. Afterwards they were placed into the measuring module of a Fourier-transform infrared spectrometer (Bruker Alpha II) in attenuated total reflection (ATR) mode. The baseline was subtracted with Origin Pro 2019 (v.9.60) and the data normalized. Additionally, crosslinking of the films with genipin and EDC/NHS was evaluated by determining the quantity of free amines using trypan blue following the method by Silva et al. [18]. The films were freeze-dried for 24 h and weighed to obtain the non-hydrated weight. Afterwards the films were rehydrated in 150 mmol NaCl (pH=4) overnight. A 0.08% trypan blue solution in 150 mmol NaCl (pH=4) was added to the films overnight at 37 °C. Afterwards the supernatants were transferred to 96 well plates and the absorbance was measured at 580 nm in a microplate reader (FLUOstar, BMG LabTech, Offenburg, Germany). A standard curve was prepared by measuring the absorbance for a series of trypan blue solutions at different concentrations. The degree of crosslinking was calculated as follows (1):

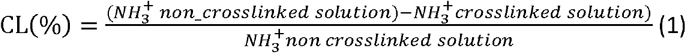

### 2.6 Oxygen permeability

Deionized water was deoxygenated by purging with N_2_ under vacuum for 15 min. This water was carefully transferred to glass bottles with a volume of 300 mL. On the bottle opening the free-standing films were fixed. Additionally, the bottles were left open (positive control) or closed with an airtight cap (blank). All bottles were constantly agitated for 24 h. Finally, the dissolved oxygen amount in the water was determined by Winkler-oximetry titration [33].

### 2.7 FGF2 upload and release

A solution of 5 µg/mL FGF2 (recombinant human FGF-basic, Peprotech, New Jersey, USA) containing 0.1 % BSA (albumin fraction V (Carl Roth GmbH + Co. KG) was prepared in PBS for loading into the films. The sterile native and crosslinked films (12 mm diameter discs) were incubated in 300µL FGF2 solution overnight (18 h was maintained for all the experiments) at 4 °C in a 24 well plate [34]. After washing the FGF2-loaded films once with PBS, 1 mL release medium (PBS, pH 7.4) was added to the film. 0.5 mL of the supernatant was collected at various time points and replaced with fresh 0.5 mL of PBS. All the supernatants were evaluated by using a human FGF-basic standard ABTS enzyme-linked immunosorbent assay (ELISA) development kit and ABTS ELISA buffer kit (both Peprotech).

### 2.8 Localization of FGF2 distribution in films

Alginate was labelled with 6-amino-fluorescein (Sigma-Aldrich Chemie GmbH, Steinheim, Germany) by EDC/NHS chemistry to obtain 10% labelled carboxylic groups, according to the protocol published by our group [35]. FGF2 was labelled with ATTO 514 NHS-Ester according to the protocol of the producer (ATTO-TEC GmbH, Siegen, Germany). Briefly, 20µg of FGF2 was dissolved in 100µL of a labeling buffer, consisting of 20 parts PBS and 1 part 0.2M sodium bicarbonate solution adjusted to pH=8.3. Afterwards, 10 µL of 5 mg/mL ATTO 514 in DMSO (Carl Roth GmbH + Co. KG, Karlsruhe, Germany) was added and the solution incubated under light protection for 1 h at RT. The resulting product was filtered with ROTI Spin MINI columns (Carl Roth GmbH + Co. KG, Karlsruhe, Germany) at 12.000 g for 10 min. The precipitate was then reconstituted in PBS and stored in Protein LoBind® Tubes (Eppendorf, Hamburg, Germany) at -20°C. Multilayer films with FL-ALG and loading with ATTO-FGF2 was done according to the procedure mentioned above. Films were transferred in 24-well cell imaging plates and a z-stack of each film with consistent settings over all samples recorded. To characterize the FGF2 distribution, the mean fluorescence intensity of ATTO-FGF2 was quantified with ImageJ (1.53c) for each z-layer.

### 2.9 Cell culture

Normal Human Dermal Fibroblasts were used in this study to investigate the metabolic activity, cytotoxicity, growth, and migration in combination with different multilayers coatings. Cryoconserved cells were thawed and cultured in Dulbecco’s modified Eagle medium supplemented with 10% fetal bovine serum 1% penicillin, streptomycin and Fungizone. HDF cells were harvested with 0.25% trypsin/0.02% EDTA solution at 37 °C for 3– 5 min. The trypsin reaction was stopped by adding DMEM with 10% FBS. Subsequently, the cells were re-suspended and seeded at a desired density on the well plate.

### 2.10 *In vitro* biocompatibility studies

The metabolic activity of HDF cells cultured exposed to the films was analysed using a resazurin assay (Deep Blue Cell Viability Kit (BioLegend, San Diego, USA)). All films were sterilized by immersion in 70% ethanol, with subsequent threefold washing with PBS for 5 min each. 1 mL cell suspension with a density of 25,000 cells per well was seeded in a 24 well plate and incubated for 12 h (DMEM + 10% FBS + 1% AB). Films were added on top of the cells after the latter were attached on the well and the medium was changed to DMEM with 1% FBS and 1% AB. After 24 h the films were temporarily transferred to a new well plate. The films were removed before each measurement, to remove effects of cell adhesion to the films. Deep Blue was prepared at a ratio of 1:10 with colourless DMEM (without pyruvate, with 4.5 g/L glucose (Lonza, Basel, Switzerland)) and was added to each existing well and incubated at 37 °C for 3 h. After incubation, duplicates of 100 µL of the supernatant were transferred to a black 96-well-plate (Greiner Bio-One International GmbH). The converted fluorescent products were photometrically quantified at an excitation wavelength of 544 nm and an emission wavelength of 590 nm using a plate reader (FLUOstar, BMG LabTech, Offenburg, Germany). Cells without films were used as negative control. After measurements were conducted, the Deep Blue medium was aspirated from the wells and was replaced by DMEM medium supplemented with 1% FBS and 1% AB pen/strep/fungi. The films, which were stored in a different well plate, were added back to the respective wells. The same procedure was repeated on day 3 and day 7.

### 2.11 Cell migration studies

Autoclaved migration fences (Aix Scientifics CRO, Aachen, Germany) were inserted into the 24 well plate and incubated for 30 min at 37 °C, to increase the attachment of the silicone sealing of the fence to the bottom of well plates. 150 µL of cell suspension containing 50,000 cells per well were seeded at the center of the migration chamber. The outer chamber was filled only with medium. This assembly was incubated for 12 h at 37 °C, 5% CO_2_. Following this, the migration chambers were removed carefully, and the wells were washed once with DPBS (with Ca and Mg (Lonza, Basel, Switzerland)). Following the removal of the migration chamber, the films were added on top of the cells and incubated for 48 h. The test medium used was DMEM + 1% FBS + 1% AB with the addition of 5 µg/mL of mitomycin C (abcr GmbH, Karlsruhe, Germany) to inhibit cell proliferation. After incubation for 24 h, the cells were washed once with PBS and fixed with 1 mL methanol Methanol (Carl Roth GmbH + Co. KG, Karlsruhe, Germany) for 10 min. Then the methanol was removed, and the cells washed with PBS. For visualization, the cells were stained with 1 mL 10% Giemsa’s azur eosin methylene blue solution (Merck KgaA, Darmstadt, Germany) in milli-Q water for 10 min. Subsequently the cells were washed once with PBS followed by ultrapure water. Series of microscopic images using 4× objective (Nikon Eclipse Ti2) were taken along the length of the diameter. These images were then stitched by using the image stitching plugin by Preibisch et al. [36](v. 1.2) which is provided as part of the Fiji project (v. 1.53c) and the length of the diameter was measured.

In an additional scratch assay the cells were seeded directly on the 24 well plate at a density of 50,000 cells per well. After incubation for 24 h, the cells were washed once with DPBS and subjected to 24 h serum starvation (DMEM without FBS). A 100 μL sterile pipette tip was used to make a uniform scratch on the well plate. The detached and floating cells were removed by washing the wells once with DPBS. 5µM CellTracker Green CMFDA (was added and was incubated for 45 minutes to stain all living cells. After washing with DPBS, medium (DMEM with 1% FBS, 1% AB and 5 μg/mL of mitomycin C) was added to the cells. At regular time points pictures were taken for 24 h with a Zeiss LSM 710 with an incubator XL S1 (Zeiss, Jena, Germany). The wound area was assessed with the help of the segmentation ImageJ plug-in Scratch Assay Analyzer developed by Glaß et al. [37]. At t=0 the wound area was set as 100%. To determine the *in vitro* wound closure rate a linear fit was applied.

### 2.12 *In vivo* biocompatibility studies

Free-standing films were implanted subcutaneously into mice to examine the tissue reaction related to the different crosslinking methods. Experimental groups were formed based on the crosslinking method used as follows: 1) native (non-crosslinked) films (N-FSF); 2) films crosslinked with EDC/NHS (E-FSF) and 3) films crosslinked with genipin (G-FSF). Prior to implantation of samples, animals were anesthetized by intraperitoneal administration of the ketamine/xylazine mixture according to the guidelines for mouse anesthesia. The skin on the back was shaved, washed with povidone iodine and incision was made. One film, 13 mm in diameter, was implanted per animal, subcutaneously, just below the interscapular region as previously published [38–40]. Each experimental group consisted of 15 animals carrying the same sample type. Implants were extracted and analyzed after 3, 10 and 30 days (five animals from each group were sacrificed per each experimental period). Extracted films with surrounding tissue were further used for histological analysis and were fixed in 10% neutral buffered formalin (NBF) until further tissue processing.

After fixation in 10% NBF, tissue samples were dehydrated by ascending concentrations of ethanol, cleared in xylene, embedded in paraffin and then sliced on a microtome Leica RM2125 RT (Leica Biosystems, Germany). The haematoxylin and eosin (H&E) and Azan trichrome (AT) staining were performed on tissue sections from five different animals per group for each experimental period and sample type. Histomorphometrical measurements were performed in NIS-Elements software version 3.2 (Nikon, Tokyo, Japan) on imaged tissue slides. The images were obtained on a microscope Leica DMLS equipped with the camera CMEX-10 Pro (Euromex Microscopen BV, Netherlands) at various magnifications. Films’ thickness after explantation and zone of cell migration into the material, on H&E and AT stained tissue sections were measured. Results are presented as mean free standing film thickness (µm) and cell migration zone (µm) ± standard deviation (SD), for each group and time point.

### 2.13 Antibacterial activity

Disc diffusion test was used to study the antibacterial activity. *E. coli* (DH5α) and *B. subtilis* DSM 10 (wild type) were the two bacteria strains used for the study. A single colony of the bacteria was inoculated in 5 mL LB liquid medium (Carl Roth GmbH + Co. KG) separately. This was then incubated overnight at 37 °C with constant agitation (160 rpm). The following day, the optical density (OD at 600 nm) was measured. The culture was diluted with sterile LB medium to obtain an OD_600_ of 1 and 200 µL of this was spread on the LB plate (using a spreader). Then, films were placed on separate quadrants of the plate. 25 µL of kanamycin (0.5 µg/µL) (Carl Roth GmbH + Co. KG) was added on sterile cellulose acetate filter paper and was used as the positive control. The negative control was sterile filter paper with autoclaved water. The plates were then incubated at 37 °C for 24 h. Later, images of the plates were taken, and the inhibition zone was measured using ImageJ (v.1.53c). Finally, the disc diameter was subtracted from the inhibition zone diameter.

### 2.14 Statistical analysis

All quantitative data were statistically processed using OriginPro 2019 (v.9.6.0.172, OriginLab Corporation). For normal distributed data (tested with Shapiro-Wilk) one-way analysis of variance (ANOVA) with a post-hoc Tukey test were used. Nonparametric data were tested with a Kruskal-Wallis test. The results of histomorphometry were statistically analyzed by (ANOVA) in SPSS software 20.0. Data are represented as mean values ± standard deviations (SD). The statistical significance is shown by asterisks in the figures (p ≤ 0.05).

## 3 Results and discussion

### 3.1 Physical characterization of free-standing films

#### 3.1.1 Studies on effects of crosslinking

The crosslinking density of FSF crosslinked by either EDC/NHS or genipin was evaluated using FTIR spectroscopy and trypan blue assay (**Figure 2A**). The FTIR-spectrum of CHI (**Figure S1**) shows characteristic, overlapping absorption bands at 1647 cm^−1^ (amide-I) and 1587 cm^−1^ (amide-II), which are the result of the 85% N-deacetylation degree of the chitin [41]. ALG is characterized by the presence of carboxylic groups, which can be found in the spectrum as carbonyl bond (C=O) at 1591 cm^−1^ [42]. In the recorded spectrum of N-FSF the overlapping absorption bands of amide-I and amide-II of CHI disappear through the electrostatic interaction with carboxylic groups of ALG [42,43]. By crosslinking with EDC/NHS and genipin, in both cases, a band shifting of the amide II band to a lower wavelength can be observed (from 1606 cm^−1^ (N-FSF) to 1602.77 cm^−1^ (E-FSF) and 1587.34 cm^−1^ (G-FSF)) (arrows in **Figure 2A**) [44]. This band shifting is characteristic for the formation of a secondary amine and indicates a higher crosslinking degree of amine groups by genipin. In the case of both crosslinked films, the N–H peak that occurred at 3293 cm^−1^ was weak and almost absent, which can be attributed to N–H being crosslinked by either genipin or EDC/NHS.

**Figure 2.**
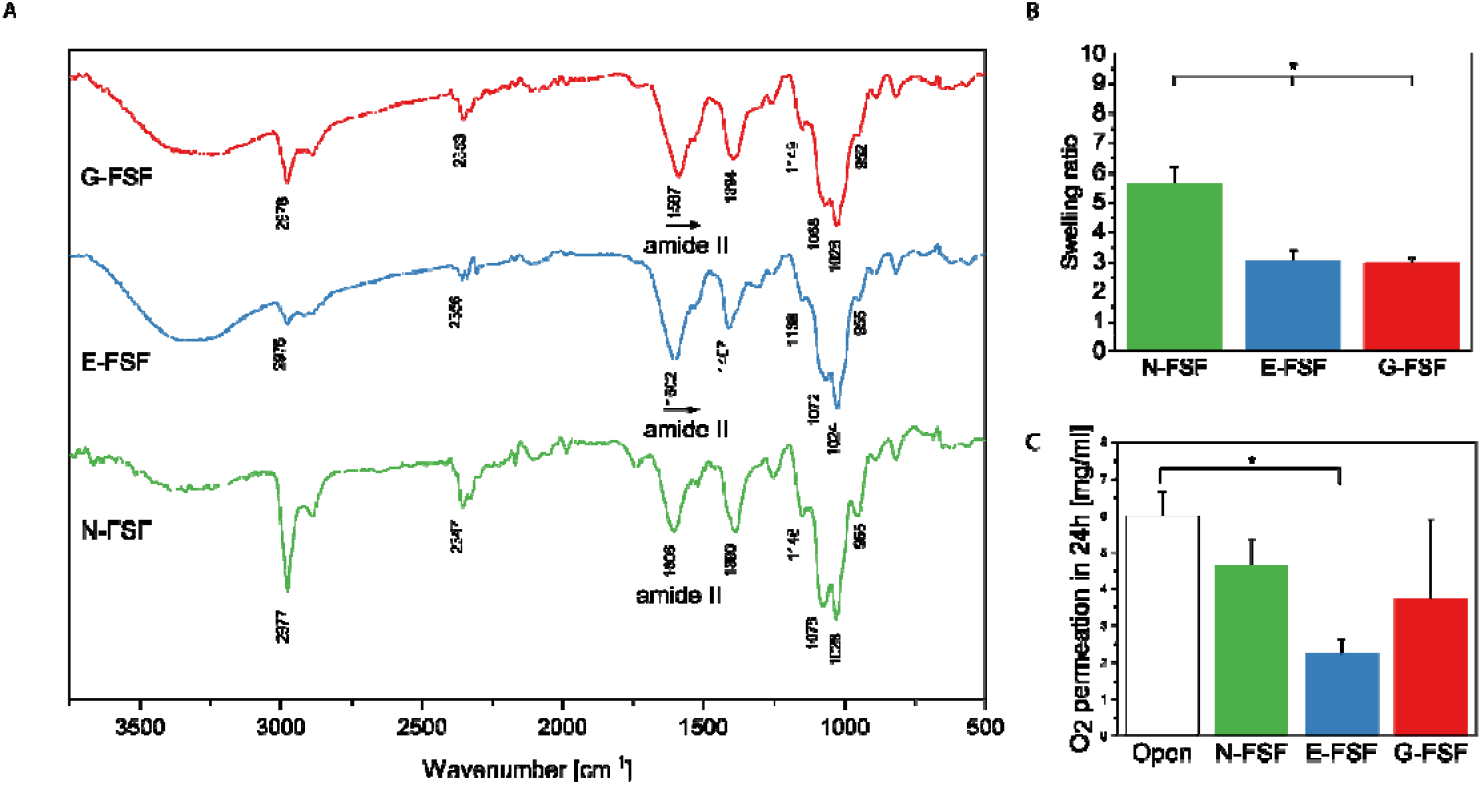
**A)** FTIR spectra of free-standing films. After crosslinking a band shift of the secondary amine groups to a lower wavelength can be observed (from 1606 cm^−1^ (N-FSF) to 1602.77 cm^−1^ (E-FSF) and 1587.34 cm^−1^ (G-FSF)). **B)** Swelling ratio determined by gravimetric method in PBS after 48 h. n=5 *p≤0.05 C) Oxygen permeation through the FSFs during 24 h. n=3 *p≤0.05

Due to the electrostatic attraction of trypan blue to free amino groups, the number of amide bonds can be assessed [45]. The calculated crosslinking degree of G-FSF of 40.6±4% is in accordance with the results of other groups employing a similar genipin crosslinking strategy for free-standing PEM films [23]. In comparison to G-FSF, the crosslinking density in the E-FSF sample was found to be significantly lower (6.9±1%). This can be explained by the different crosslinking chemistry of EDC/NHS compared to genipin (**Figure 1**). EDC/NHS forms covalent amide bonds between amino groups of CHI and carboxylic groups of ALG. Through this, all sterically accessible CHI and ALGs, are crosslinked. Genipin on the other hand crosslinks amino groups only and therefore is only capable of crosslinking CHI molecules. Then, ALG is only connected to CHI by ion pairing with the remaining amino groups of CHI. Indeed, the presence of genipin was validated by the very broad fluorescence of G-FSF [46], which was used to ensure batch-to-batch reproducibility (**Figure S2**).

#### 3.1.2 Oxygen permeability

The oxygen permeability of FSF is of interest due to their anticipated application as wound dressing that should enable a sufficient oxygen supply to the damaged tissue [47]. Moreover, oxygen permeability is linked to angiogenesis and cell proliferation, inhibits growth of anaerobic bacteria and enhances leukocyte bacterial killing capacity [48,49]. E-FSF possessed the least oxygen permeability compared to G-FSF and N-FSF (**Figure 2B**). Although none of the films reaches an oxygen permeation like the open control, the N-FSF and G-FSF showed a non-significant reduction, which in case of the G-FSF was similar to results of other studies [23]. The degree of swelling shown in **Figure 2B** is related positively with the oxygen permeability. This might be caused by the swelling of the polymer chains, which increases the O_2_ penetration as well as the transport of O_2_ through the water present in the films [50].

#### 3.1.3 FGF2 uptake and release

Bioactive wound dressing should support neovascularization and promote regeneration of skin. This can be promoted by the growth factor FGF2 which represents a strong mitogen, promoting growth of fibroblasts, endothelial and other cells [8]. The amount and distribution of uploaded fluorescence labelled FGF2 (ATTO-FGF2) was studied by CLSM z-stack quantification. In all films ATTO-FGF2 was present in the core of the film, confirming diffusion throughout the layers (**Figure3A and S3**). FGF2 is known to interact with the amino groups of CHI which protects the growth factor against denaturation by heat, acidic pH, and proteolysis [51]. At the same time the basic epitopes of FGF2 can bind to negatively charged carboxylic groups of ALG [52]. The total uptake of FGF2 into the N-FSF was higher than in G-FSF, which was higher compared to E-FSF. This is indicated by the distribution of the ATTO-FGF2 3D stacks in the FSF (**Figure 3B**). Accordingly, a higher cumulative release of FGF2 from N-FSF (97±5 ng/mL) was measured after 7 days in comparison to the crosslinked films (E-FSF: 19±3 ng/mL and G-FSF: 39±1 ng/mL) by FGF2 ELISA (**Figure 3C**). The high uptake of N-FSF can be accounted to the higher swelling ratio, which increases absorption of FGF2. Additionally, the release from N-FSF is faster, caused by the easier diffusion of FGF2 through the films probably because of the looser packing of the polyelectrolytes and comparable weaker electrostatic interactions between CHI and ALG at pH 7.4.

**Figure 3.**
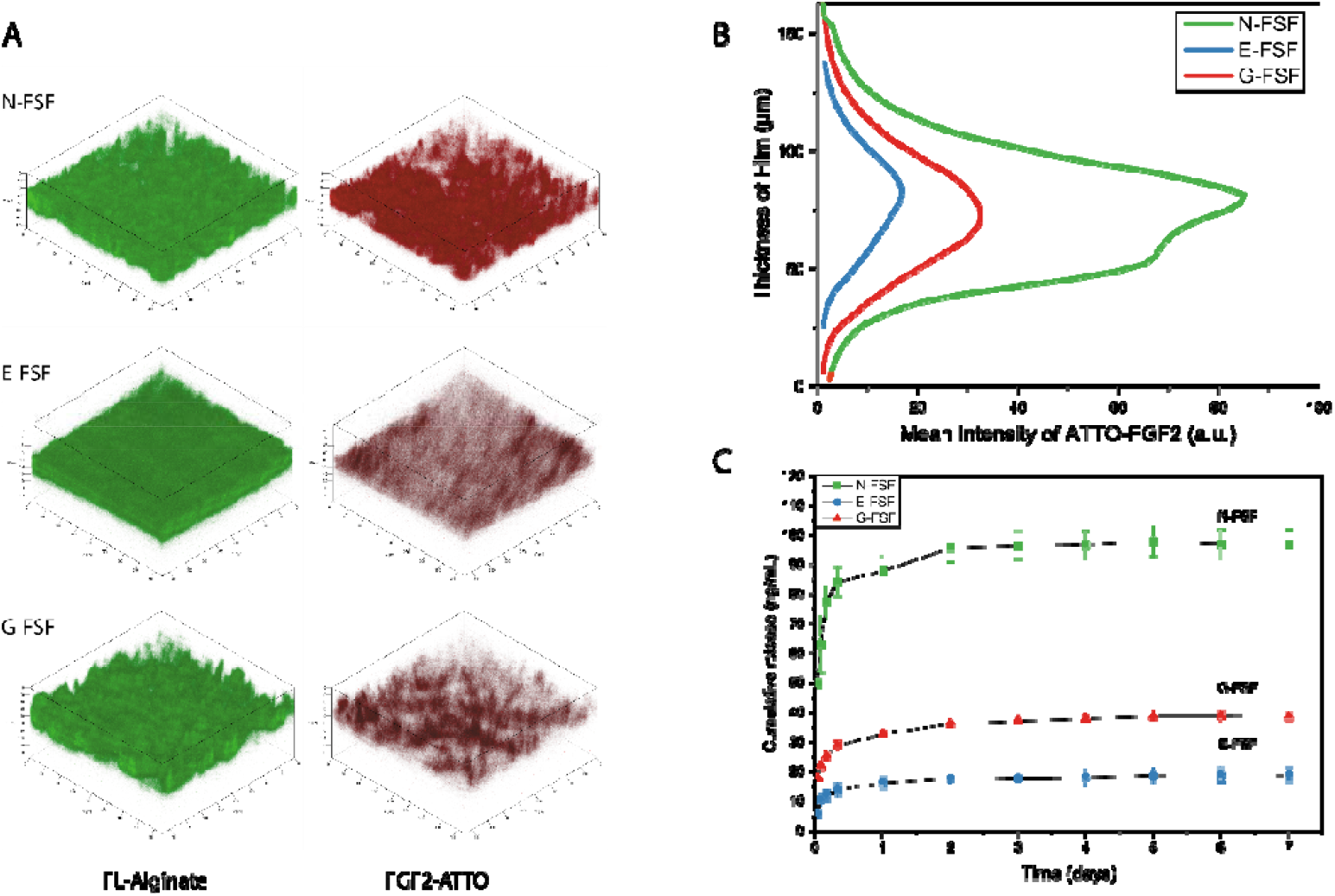
**A)** 3D-volume renderings based on z-stacks of films recorded by CLSM to determine the distribution of FGF2 in the films. Green is FL-ALG. Red is FGF2-ATTO. **B)** Mean Intensity of FGF2 in each z-layer throughout the thickness of the fluorescent-labelled film. Area under Curve of each condition is significantly different. (n=6, *p≤0.05) **C)** Cumulative release of FGF2 from the films over the duration of 7 days. Half of the release medium was discarded, to emulate sink conditions in part. The release was quantified by ELISA. The slopes are all significantly different to each other. (n=3, *p≤0.05)

It is important to note that the initial burst from N-FSF (49.7±2 ng/mL) was significantly higher than from the crosslinked films (E-FSF: 6.0±1 ng/mL and G-FSF: 17.9±0.11 ng/mL). The release slopes were statistically significant from each other (N-FSF: 6.2 ng/d, E-FSF: 1.9 ng/d and G-FSF: 3.9 ng/d) and were fitted to different models of release shown in the (**Table S4**). For both crosslinked and native films the correlation coefficient (r^2^) for the Higuchi model was the highest, indicating that the release is determined by the diffusion through polymers [53]. These findings are similar to studies of FGF2 released from poly (methacrylic acid)/poly-l-histidine polyelectrolyte multilayers [54]. In the case of crosslinked FSF, a dense network created by crosslinking is known to decrease and delay the release [55,56]. However, even though the crosslinking density of G-FSF is higher than of E-FSF it was able to take up more FGF2 with higher cumulative release. This might be related to the presence of non-crosslinked ALG, which remaining carboxylic groups can undergo ionic interactions with the basic epitopes of FGF2 [56]. Additionally, the presence of genipin with its indol alkaloid backbone might lead to the association of aromatic amino acids [57]. This may explain the occurrence of high amounts of bound FGF2.

### 3.2 Biological characterization

#### 3.2.1 *In vitro* biocompatibility studies

To assess the biocompatibility and the mitogenic effect of the films, with and without FGF2-loading, a growth assay with HDF cells was conducted. The number of cells directly attached to the films was found to be very low (**Figure S5**), which can be regarded as a positive property of a wound dressing [2].Therefore, in all further experiments cells cultured on tissue culture plastic (TCP) were evaluated, excluding cells adhering to the films. As seen from **Figure 4**, the cell viability for G-FSF on day 1 was lower compared to that of the control (cells without films). G-FSF on day 1 caused a lower cell growth compared to all other FSF (N-FSF and E-FSF). However, on day 3 and day 7 the same G-FSF samples stimulated the growth of cells compared to the control and E-FSF. This statement can be extended to all films as the cell viability of cells subjected to the different FSF was significantly higher compared to the respective controls on day 3 and 7. The cultured fibroblast can interact with the highly deacetylated CHI (85%), which is known to enhance the proliferation, by interaction with serum proteins [58]. This effect might be supported by the ability of G-FSF to delay the FGF2 release, leading to higher levels of active FGF2 from day 3 of the experiment on. Overall, N-FSF and the crosslinked films (E-FSF and G-FSF) did not exhibit any cytotoxicity and an increase in cell proliferation was verified for all tested films.

**Figure 4.**
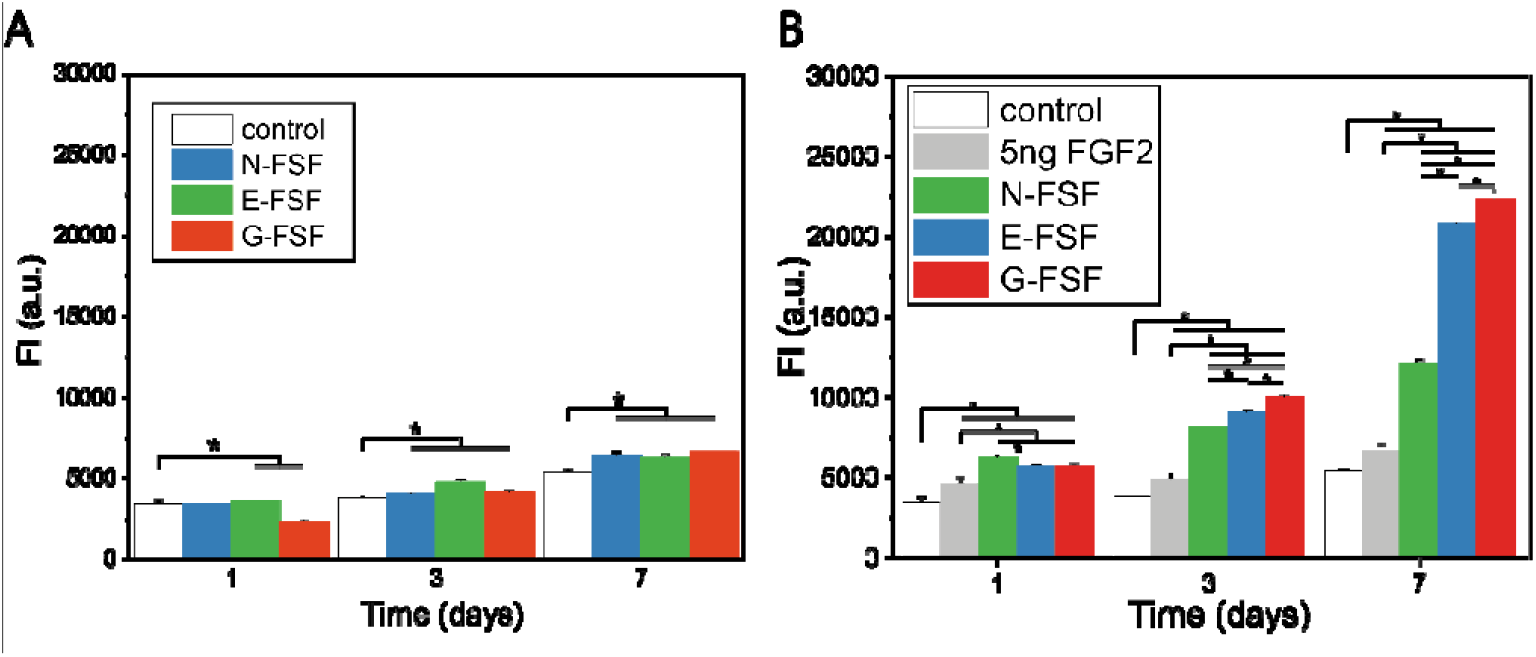
**A)** Biocompatibility testing of films towards fibroblasts (HDF) determined by Deep blue assay. The intensity values are a measure of the metabolic activity. Cells were grown on tissue culture plastic (TCP) with the films above. DMEM Medium was changed on day 1 and 3. (n=6, *p≤0.05). **B)** Cell growth assay with FGF2 loaded FSF. Experimental setup as described in A

The effect of FGF2 loading on fibroblasts growth was studied in the same manner. It was found that the presence of FGF2 in the films stimulates the growth of the cells significantly. After day 3 all FGF2 loaded films showed significantly more cell growth than the control. After day 7 G-FSF promoted the highest cell growth closely followed by E-FSF and with a certain distance N-FSF. Reasons for the stronger promotion of fibroblast growth by crosslinked films are related to the differences in the concentration and kinetics of FGF-2 release. The proliferative effect of FGF2 is concentration dependent; Low concentrations of FGF2 (<1 ng/mL) and high concentrations of FGF2 (100 ng/mL) are known to trigger survival and differentiation while delaying or inhibiting proliferation. Intermediate concentrations (1-10 ng/mL) are stimulating cell proliferation [59]. Another important aspect is the low stability of FGF2 in solution, particularly in presence of cells where it is proteolytically degraded within 24 as shown previously [14]. Hence, a fast release of larger quantities of FGF2 at early stages of cell growth will be less effective than a slower release over time. We suggest that the FGF2 loaded crosslinked films (E-FSF and G-FSF), kept the concentration of FGF2 in cell growth promoting range (1-10 ng/mL) [56,59]. FGF2 loaded N-FSF showed a comparatively high burst release (97.01 ng/mL), which was in the growth inhibitory range of FGF2. Additionally, FGF2, being positively charged at pH 7.4, can bind to the ALG layer and be presented to the cells as matrix-bound FGF2, which allows prolonged presentation of the growth factor to the cells [56]. This possibly allows G-FSF, with its non-crosslinked ALG, to increase the cell growth further. Overall, it can be stated that FGF2 loaded crosslinked FSF significantly increased the cell growth of fibroblasts compared to N-FSF, while utilizing lower amounts of FGF2.

Migration of fibroblasts and other cells plays an important role in wound healing to restore the damaged tissue. Hence, we studied here migration of fibroblast with two different methods. In both assays Mitomycin C, as a mitotic inhibitor, was added to inhibit changes in the positioning of cells due to expansion of the growing cell layer. As seen in **Figure 5A**, the films loaded with FGF2 showed a significantly larger diameter than all non-loaded films. FGF2 loaded G-FSF showed the highest cell migration, followed by E-FSF, while N-FSF exhibited the lowest migration of fibroblasts. A comparable behavior was observed in an *in vitro* scratch assay, which mimics cell migration during wound healing *in vivo* more closely.

**Figure 5.**
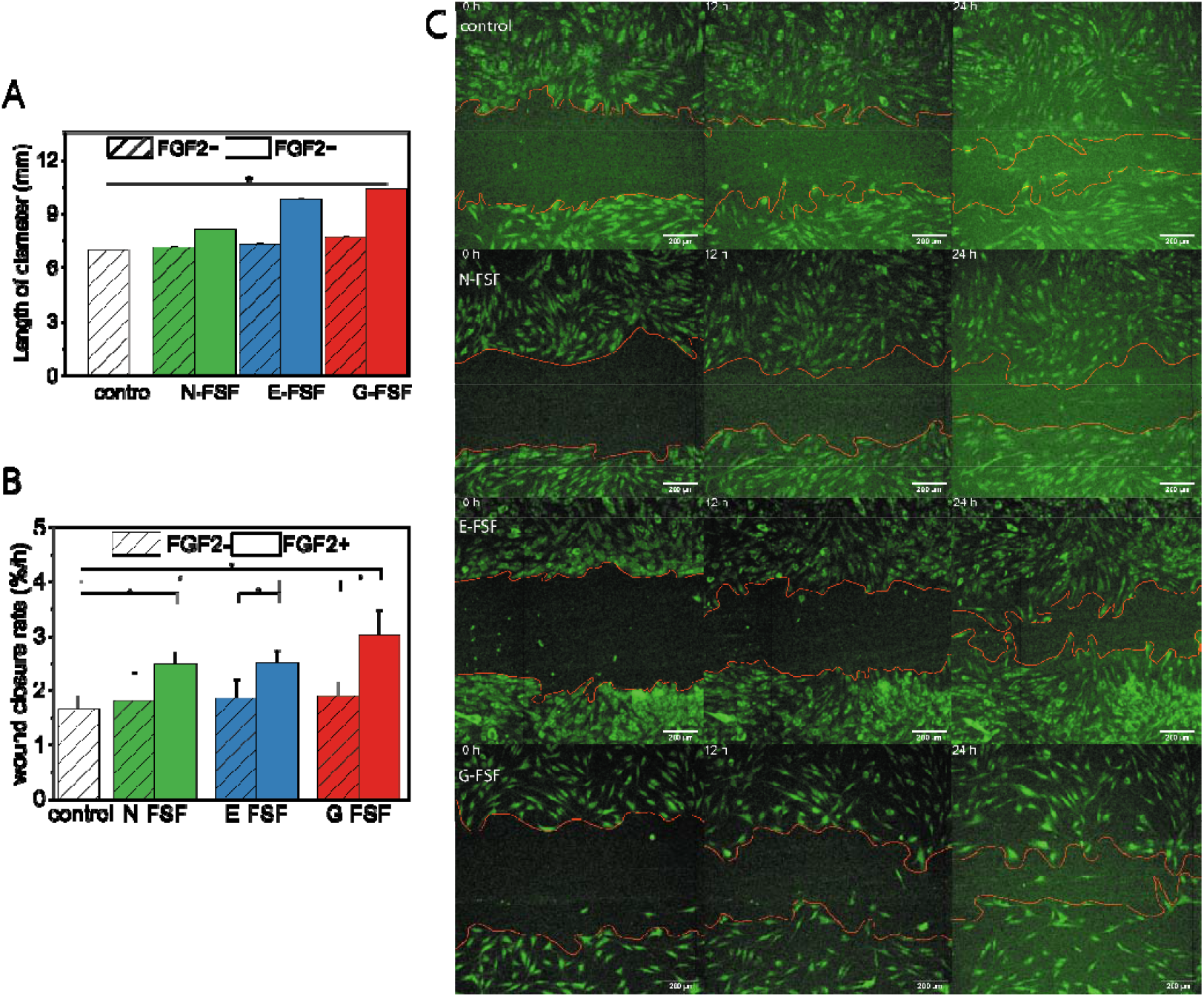
**A)** Cell migration after 48h underneath FGF2-loaded and non-loaded films on TCP. Control represents cells without any films. A larger diameter corresponds to a faster cell migration. (n=6, *p≤0.05). **B)** Linear fit of cell free area every 1 h over 24 h during an in vitro scratch assay. The addition of FGF2 leads to a significant acceleration of wound healing compared to the control. (n=6, *p≤0.05). **C)** Micrographs at selected time points of in vitro scratch assay also found as time stack **Video S7**. The area between the red lines at t=0 was set as 100%. Scale bar = 100 µm. All conditions are with FGF2 (except control: 1% FBS). (n=4 *p≤0.05)

Here, the cell-free area was determined over 24 h and subsequently the *in vitro* wound area closure rate calculated (**Figure 5B**). The G-FSF loaded with FGF2 showed the fastest *in vitro* wound closure (**Figure 5C**). Followed by E-FSF and N-FSF. Remarkable is the similarity in the initial decrease of wound area for all conditions. However, G-FSF and E-FSF permitted a faster *in vitro* wound closure over the whole duration of 24 h. These results confirm the positive effects of FGF2 as stimulator of migration in a possible wound environment [9]. The incorporation of FGF2 into the films leads to a faster migration and *in vitro* wound closure. The crosslinking with genipin seems to modulate the burst release of FGF2 in a way, which is beneficial for the migration of cells. The higher concentration of FGF2 in the medium, released in 24 h by N-FSF (88±5 ng/mL) seem to be inferior to the medium concentration of G-FSF (33±1 ng/mL) or E-FSF (16±2 ng/mL). As reported in the literature around 25 ng/mL FGF2 concentration is an optimum for migration and *in vitro* wound healing [60,61].

#### 3.2.2 *In vivo* biocompatibility

To extend the biocompatibility study to living organisms, soft tissue reaction to FSF was examined in a subcutaneous implantation model in mice. Samples were taken at 3, 10 and 30 days after implantation. Macroscopic analysis revealed that there was difference in reaction of surrounding tissue to the films (**Figure 6**). For N-FSF a significant change in shape can be seen from the 3^rd^ to the 30^th^ day of implantation in terms of degradation and loss of integrity (**Figure 6-a, d, g**). No noticeable adverse tissue reaction was observed. E-FSF induced a notable surrounding tissue reaction directly from the onset of implantation. A very pronounced surrounding tissue vascularization, swelling and granulation could be observed, which are signs of inflammatory tissue reaction to the foreign material (**Figure 6-b, e, h**). On the other hand, G-FSF was the most stable over time and did not induce visible macroscopic signs of adverse reactions after longer time post implantation (**Figure 6-c, f, i**).

**Figure 6.**
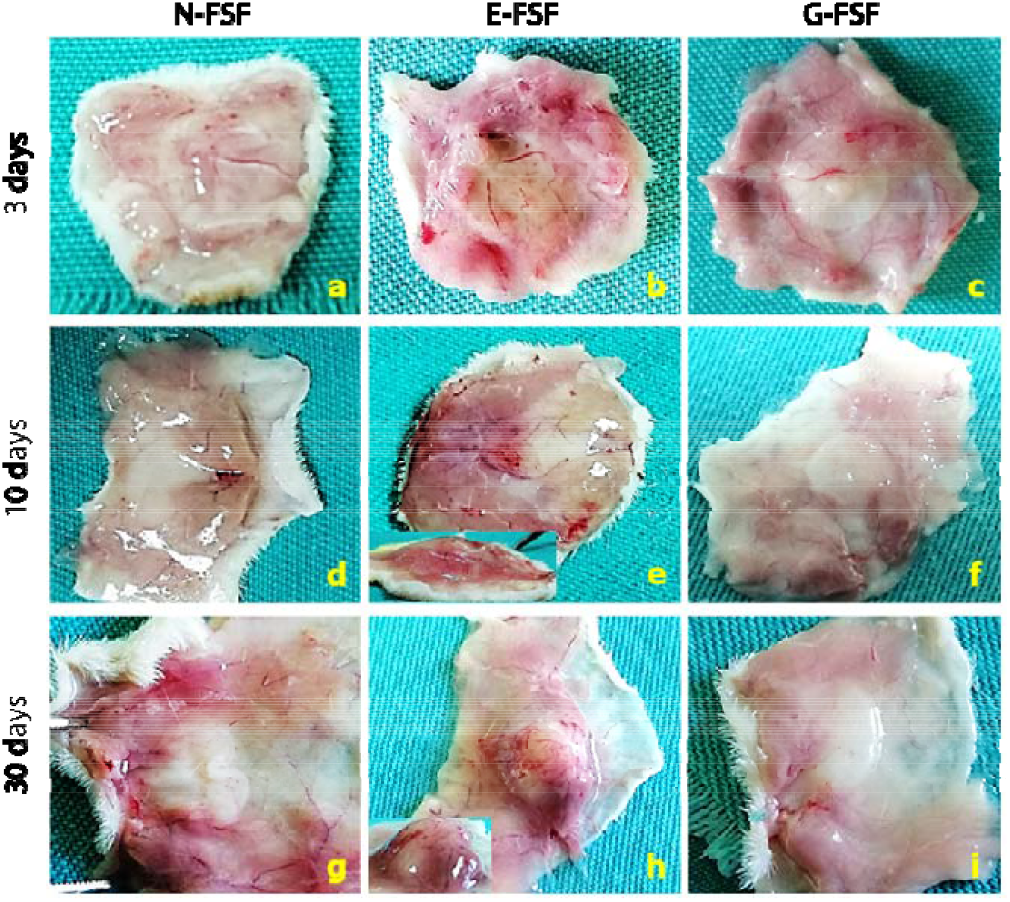
Macroscopic appearance of explanted free-standing films and surrounding tissue reaction 3, 10 and 30 days after implantation

Histological analysis was undertaken to further study the host reaction to implantation of FSF (**Figure 7**). After 3 days, an intense cell infiltration was observed in all groups. However, the presence and arrangement of cell populations differed. In the N-FSF group connective tissue cells from surrounding tissue could be seen, which aligned onto the films, went inside the films and started degradation. Additionally, signs of stratification as well as induction of collagen synthesis were observed. In the E-FSF and G-FSF group, infiltration of inflammation-related cells, such as neutrophils and macrophages, appeared in the surrounding tissue. Further, increased vascularization of surrounding tissue was noticed for both E-FSF and G-FSF. This suggests the initiation of an inflammatory reaction to the implanted films, which is an expected reaction to implanted biomaterials at that time point.

**Figure 7.**
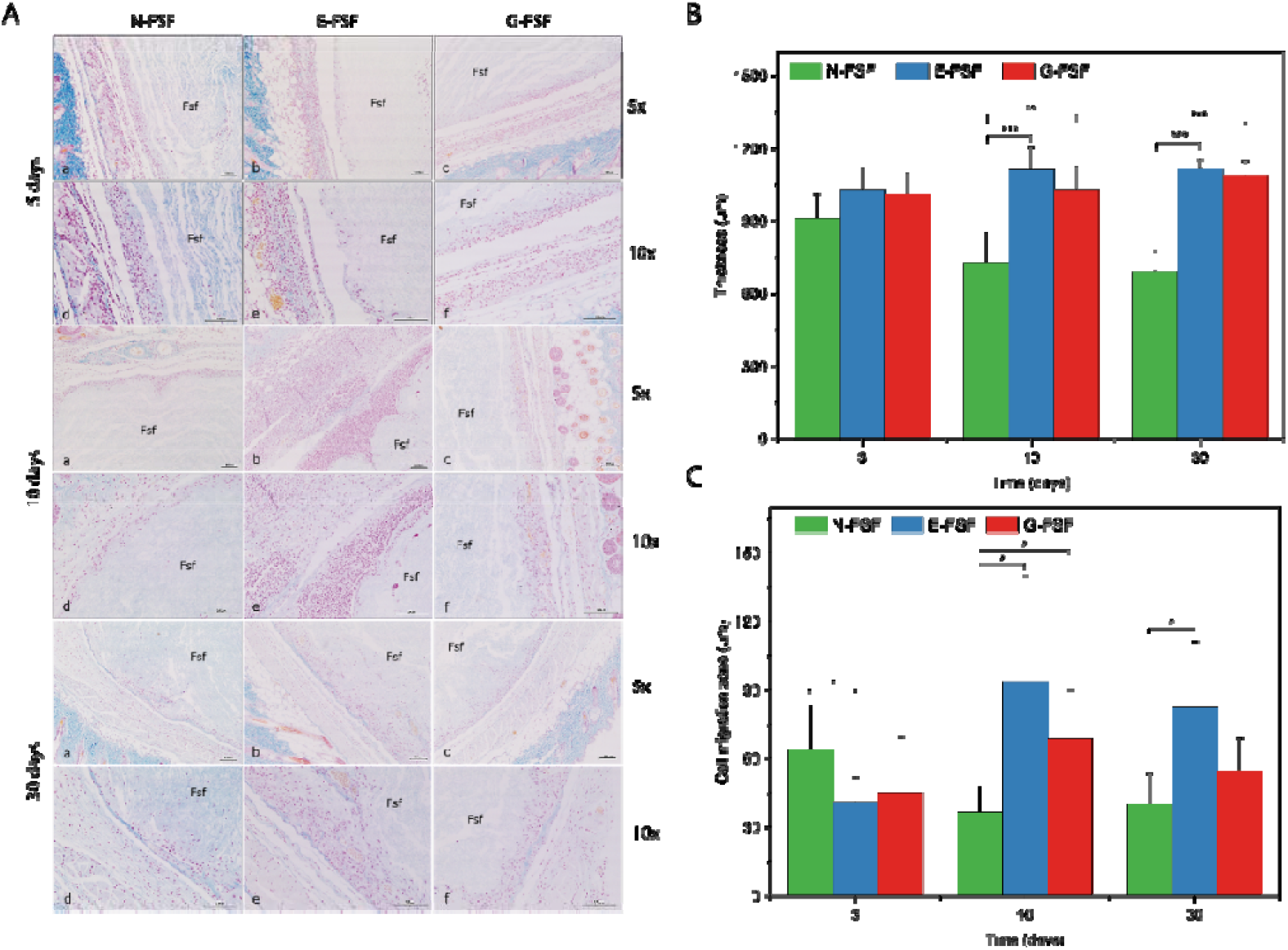
**A)** Histologic sections of explanted free-standing films with surrounding tissue 3, 10 and 30 days after implantation; AT staining; brightfield; objective magnification 5x (a-c) and 10x (d-f); blue colour on the images indicates collagen staining; Fsf – free-standing films; scale bar = 100 μm. **B)** Histomorphometric measurements; thickness (μm) of explanted free-standing films 3, 10 and 30 days after implantation and **C** cell migration zone (μm) in explanted free-standing films 3, 10 and 30 days after implantation (*) p≤0.05, (**) p≤0.01, (***) p≤0.001; n=5 for each group

After 10 days, the inflammatory tissue reaction was reduced for N-FSF and G-FSF-groups, but not in the E-FSF group, which exhibited a strong inflammatory response. In the E-FSF group large infiltrates of inflammatory cells and connective tissue cells, which are lined onto the films and started migrating inside the film could be found. Additionally, in the surrounding tissue blood vessels were found in greater number, compared to the N-FSF and G-FSF group. EDC/NHS has shown toxic effects in in-vivo studies, where it was used in comparable concentrations [62]. While the EDC/NHS crosslinking method usually yields biocompatible systems, at higher concentrations cytotoxic EDC and reaction by-products cannot be fully removed via simple rinsing [63]. N-FSF seemed to be loosely packed, with great number of connective tissue infiltrates, but without visible signs of inflammatory reaction at this time point.

At day 30, in the N-FSF group, single cells migrating again deeper into the pores of the partially degraded N-FSF as well as newly formed collagen fibers and blood vessels surrounding the films could be observed. The inflammatory reaction seen in the E-FSF group at day 10 seemed to be reduced at day 30, possibly showing the elimination of cytotoxic compounds in the E-FSF group. A lot of cells migrated into the films and deposited collagen, but also blood vessels surrounding and inside the films can be observed. This could be the consequence of strong inflammatory tissue reaction in earlier time points of the E-FSF group. In the G-FSF group no signs of inflammatory reaction of the surrounding tissue could be observed. The connective tissue cells were well integrated in the films depositing collagen fibers onto the films.

Histomorphometric measurements indicated that indeed the thickness of the N-FSF (**Figure 7B**) gradually decreased from day 3 to day 30, which suggests *in vivo* degradation. Crosslinked FSF were significantly more stable against degradation compared to N-FSF. Significant difference in cell infiltration and migration into implanted films from day 3 to day 30 was noticed for N-FSF and E-FSF group (**Figure 7C**). N-FSF-groups cell migration zones decreased over time, while showing initially the thickest migration zone. An increase in cell infiltration and migration at day 10 was observed in E-FSF and G-FSF group with a slight decrease at day 30. Overall, N-FSF and G-FSF lead to a controlled tissue response towards their implantation, while N-FSF was showing degradation after 10 days compared to the more stable G-FSF but both FSF represent obviously biocompatible materials in vivo. By contrast, E-FSF showed a pro-inflammatory effect towards the surrounding tissue which makes this crosslinking method less suitable for making of FSF in wound dressing applications.

### 3.2.3 Antibacterial activity

Wound dressings should have also an anti-bacterial activity because infection is a common occurrence in chronic wounds [6]. Here, the antibacterial activity of FSF was studied with *E. coli* DSM 6897 as Gram-negative (G-) and *B. subtilis* DSM 10 as Gram-positive (G+) germs using the disc diffusion method (**Figure 8A**). These bacteria are widely used to study the antibacterial activity of wound dressings [64]. The positive control (12 mm filter paper disc impregnated with kanamycin produced a large zone of inhibition for both *E. coli* and *B. subtilis* as visible in **Figure 8B**. In general, the inhibitory effect of all FSF (cut in 12 mm discs) is lower compared to the aminoglycoside antibiotic kanamycin used as a positive control. It is visible that the native FSF shows a comparable effect on *E. coli* like kanamycin but lower for *B. subtilis*. Both crosslinked films show a decreased antibacterial effect indicating that a lowered release/mobility of CHI is probably responsible for that. Indeed, CHI is most probably responsible for the antibacterial effect as shown in other studies [31,65]. In summary, all FSF possess antibacterial activity which is an important finding that further qualifies the FSF developed here for a future application as wound dressing.

**Figure 8.**
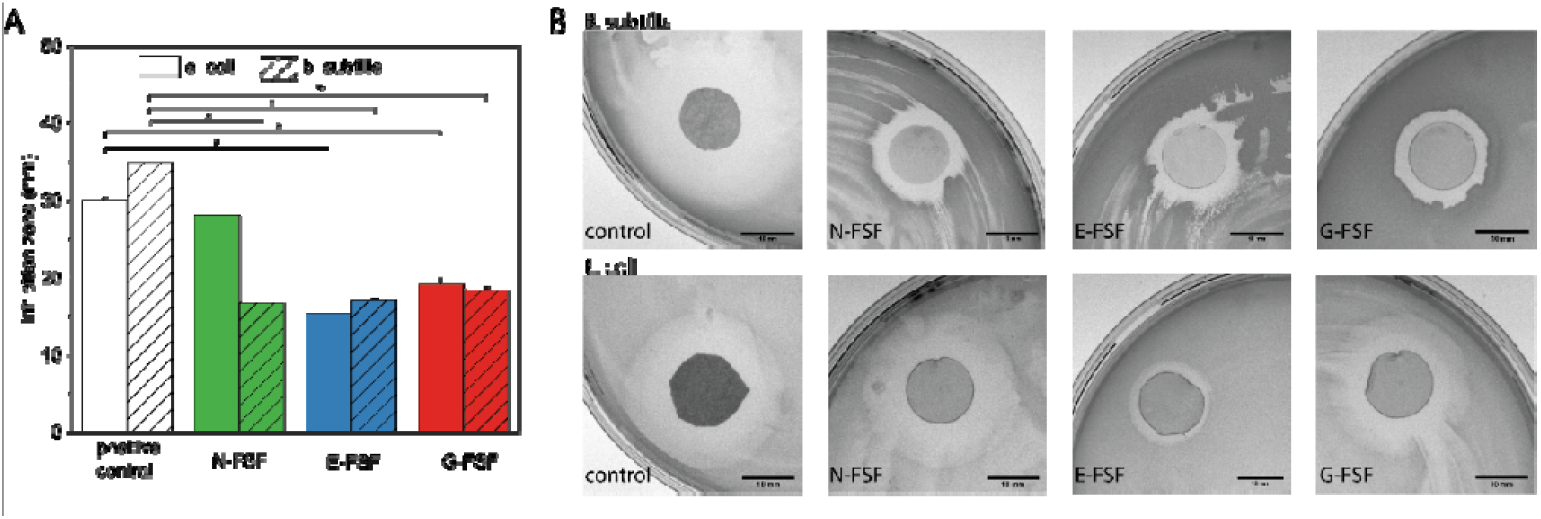
**A)** Antibacterial activity of films towards *E. coli* DSM 6897 (G-) and *B. subtilis* DSM 10 (G+). Positive control is kanamycin (0.5 g/L). (n=4, *p≤0.05**). B)** Micrographs of antibacterial test. Scale bar = 10mm.

## 4 Conclusions

Here, we could demonstrate that the use of LbL-based dip coating enables the design of FSF, based on the cost effective combination of chitosan, as an antibacterial agent and alginate, which provides swelling capabilities useful for uptake of wound exudates. The most interesting finding are as follows; Additional crosslinking with genipin increases the stability of films *in vivo* and allows the control of FGF2 release, and, in notable contrast to N-FSF, it also improves the general biocompatibility *in vitro* and *in vivo* with a promoting effect on migration and growth of fibroblasts. Hence, free-standing films prepared by LbL-process have a great potential to serve as wound dressing materials that may support regeneration of dermis as a prerequisite for the re-epithelization of wounds. All together the proposed FSF made of alginate and chitosan, crosslinked with genipin and loaded with FGF2 may hold great promises for use as wound dressings which will be studied in future investigations.

## Abbreviations

ALG: alginate
CHI: chitosan
EDC: 1-ethyl-3-(−3-dimethylaminopropyl) carbodiimide
E-FSF: EDC/NHS crosslinked free-standing film
ELISA: enzyme-linked immunosorbent assay
FGF2: fibroblast growth factor 2
FL-ALG: fluorescein labelled alginate
FSF: free-standing films
GF: growth factor
G-FSF: genipin crosslinked free-standing film
HDF: human dermal fibroblasts
LbL: layer-by-layer
N-FSF: native free-standing film
NHS: *N*-hydroxysuccinimide
PEM: polyelectrolyte multilayer

## Author contributions

**Adrian Hautmann:** Conceptualization, Methodology, Software, Investigation, Writing - Original draft, Visualization **Devaki Kedilaya**: Conceptualization, Methodology, Investigation, **Sanja Stojanović:** Methodology, Investigation, Writing - Review & Editing **Milena Radenković:** Methodology, Investigation **Christian K. Marx:** Methodology, Investigation **Stevo Najman:** Resources, Writing - Review & Editing, Supervision **Markus Pietzsch:** Resources, Writing - Review & Editing, Supervision. **João F. Mano:** Conceptualization, Writing - Review & Editing **Thomas Groth:** Conceptualization, Methodology, Resources, Writing - Reviewing and Editing, Supervision, Project administration, Funding acquisition

## Acknowledgments

The International Graduate School AGRIPOLY funded by the European Regional Development Fund (ERDF) and the Federal State Saxony-Anhalt supported this work. Additionally, this project was supported by a Project-Related Personal Exchange program funded by the Ministry of Education, Science and Technological Development of the Republic of Serbia and the German Academic Exchange Service (DAAD).

