## Supplemental information for "Free-Standing Multilayer Films as Growth Factor Reservoirs for Future Wound Dressing Applications"

*[1]Department of Biomedical Materials, Martin Luther University Halle-Wittenberg, Germany, [2]Department for Cell and Tissue Engineering, Scientific Research Center for Biomedicine, Faculty of Medicine, University of Niš, Niš, Serbia, [3]Department of Biology and Human Genetics, Faculty of Medicine, University of Niš, Niš, Serbia, [4]Department of Downstream Processing, Institute of Pharmacy, Martin Luther University Halle-Wittenberg, Germany, [5]CICECO—Aveiro Institute of Materials, Univeristy of Aveiro, Protugal, [6]Interdisciplinary Center of Material Research, Martin Luther University Halle-Wittenberg, Germany*

A. Hautmann, D. Kedilaya, T. Groth
Department of Biomedical Materials, Martin Luther University Halle-Wittenberg
Heinrich-Damerow-Strasse 4, 06120, Halle (Saale), Germany

C. Marx, M. Pietzsch
Downstream Processing, Institute of Pharmacy, Martin Luther University Halle-Wittenberg
Weinbergweg 22, 06120 Halle (Saale)

J.F. Mano CICECO—Aveiro Institute of Materials, Department of Chemistry, University of Aveiro, 3810-193 Aveiro, Portugal

T. Groth
Interdisciplinary Center of Material Science, Martin Luther University Halle-Wittenberg, 06099, Halle (Saale), Germany

### Supplement

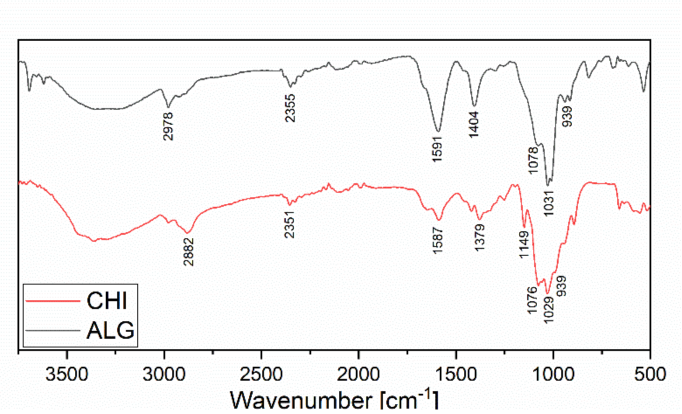


S 1 – FTIR spectra of chitosan (85% deacetylated) and alginate.

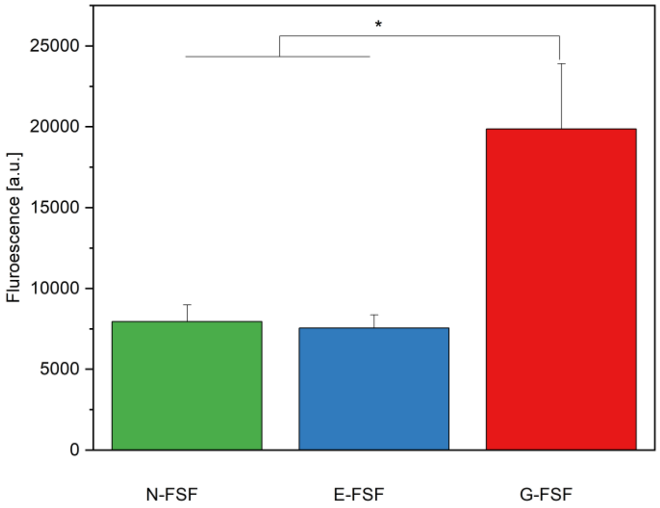


S 2 - Fluorescence of films. Fluorescence of genipin confirms the successful crosslinking.

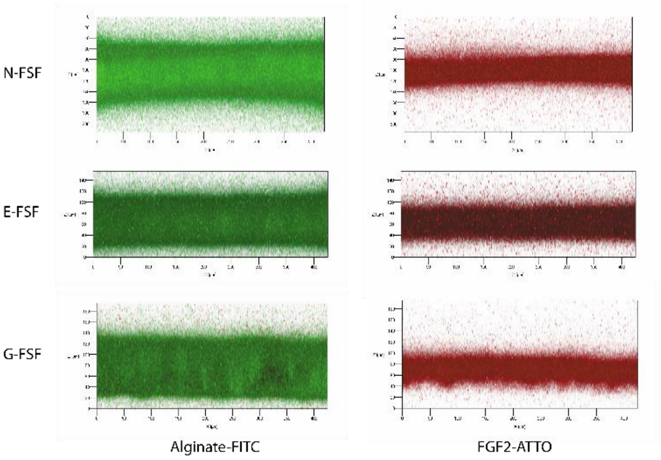


S 3 – Z-stack side view of free-standing films. Green is ALG-FITC red is FGF2-ATTO

| Model Name | Non-crosslinked | | | EDC-NHS | | | Genipin | | |
|---|---|---|---|---|---|---|---|---|---|
| | $r^2$ | Slope | Intercept | $r^2$ | Slope | Intercept | $r^2$ | Slope | Intercept |
| Zero order model | 0.7827 | 17.134 | 16.758 | 0.8315 | 13.99 | 9.9938 | 0.7209 | 13.805 | 15.952 |
| First order model | 0.9262 | -0.1465 | 1.9233 | 0.8904 | -0.094 | 1.9555 | 0.822 | -0.0972 | 1.9165 |
| Hixson-Crowell model | 0.8822 | 0.4091 | 0.2769 | 0.8721 | 0.2876 | 0.1592 | 0.7877 | 0.2922 | 0.2806 |
| Korsmeyer-Peppas model | 0.6051 | 89.267 | 26.588 | 0.7253 | 77.427 | 16.996 | 0.5364 | 70.56 | 24.18 |
| **Higuchi model** | **0.9713** | **38.698** | **4.0368** | **0.9671** | **30.591** | **0.7188** | **0.9426** | **32.004** | **4.7922** |

S 4 - Correlation of release models for the FGF2 release out of films in terms of slope, intercept and correlation coefficient ($r^2$).

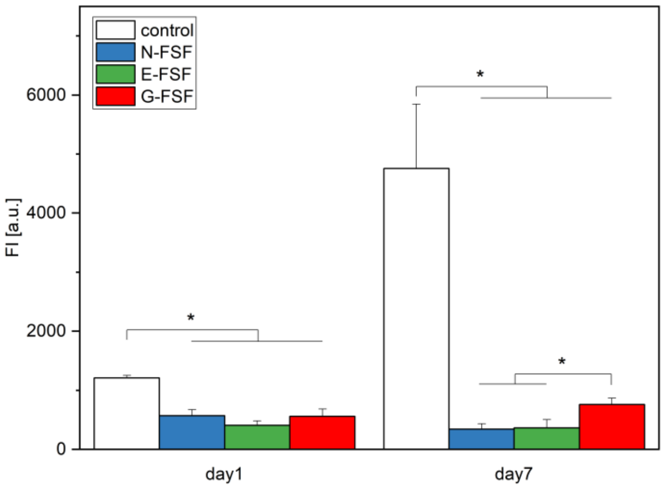


S 5 – Cell viability of cells only adherent directly to the film.

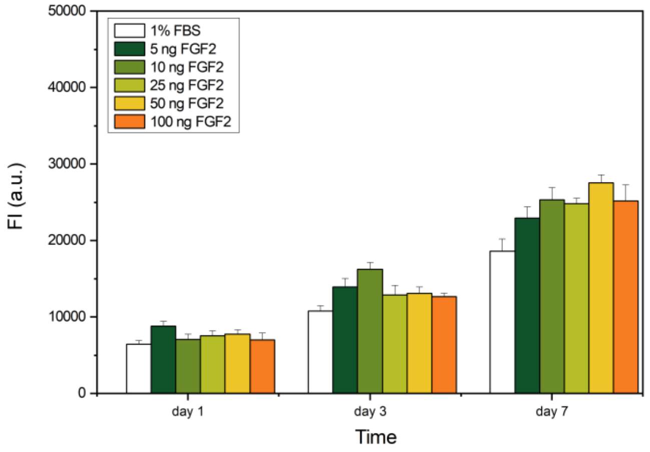


S 6 – Qblue Assay on HDF cells grown on TCPS to assess the optimal amount of FGF2 amounts in terms of cell growth.

S 7 – Time stacks - The area between the red lines at t=0 was set as 100%. Every 60min the area was measured. Scale bar is 100µm. All conditions are with FGF2 (except control: 1% FBS). n=4 n=3 *$p \leq 0.05$
